# Endogenous nucleophilic scavengers of reactive acyl-species neutralize carbon stress

**DOI:** 10.64898/2026.04.06.716549

**Authors:** Min-Keun Lee, Seon Min Kim, Sanghyuk Lee, Jin Ho Yang, Chang Ryul Choi, Ting Miao, Jong-Seo Kim, Norbert Perrimon, Yun Pyo Kang, Hyun-Woo Rhee

## Abstract

Aberrant protein acylation by reactive acyl species (RAS)—termed carbon stress—is a major driver of aging and metabolic disease. While enzymatic deacylases such as sirtuins counteract aberrant acylation, whether endogenous metabolites can directly neutralize RAS remains unclear. Here, we report that nucleophilic metabolites—taurine, spermidine, and ethanolamine—react with acyl-CoAs, preventing aberrant protein acylation and potentially extending lifespan. We demonstrate that spermidine scavenges acetyl-CoA within the catalytic pocket of p300, a histone acetyltransferase, and extends lifespan in *Drosophila*. Taurine supplementation in mice fed a high-fat diet promotes N-fatty acyl taurine formation, confirming in vivo scavenging of RAS. These findings identify endogenous nucleophilic metabolites as scavengers that neutralize carbon stress, with implications for combating aging and metabolic disease.

**Summary:** Protein homeostasis is constantly challenged by reactive metabolic intermediates that drive aberrant post-translational modifications, contributing to aging and metabolic disease. While the antioxidant systems that neutralize reactive oxygen species (ROS) are well characterized, whether analogous defenses exist against reactive acyl species (RAS)—the metabolic drivers of non-enzymatic protein acylation, termed carbon stress—has remained unknown. Here we show that three abundant endogenous nucleophilic metabolites—taurine, spermidine, and ethanolamine—directly intercept reactive acyl-CoA species through nucleophilic attack, preventing aberrant protein acylation. We demonstrate that spermidine scavenges acetyl-CoA within the catalytic pocket of the histone acetyltransferase p300, effectively buffering p300-mediated hyperacetylation of muscle, cytosolic, and mitochondrial proteins and extending lifespan in a Drosophila model of p300-driven toxicity. In mice fed a high-fat diet, taurine supplementation promotes the formation of N-fatty acyl taurines, providing direct in vivo evidence for nucleophilic scavenging of diet-derived RAS. Our findings reveal a previously unrecognized metabolite-based defense system—parallel to classical antioxidant defenses against ROS—that neutralizes carbon stress, with broad implications for understanding the molecular mechanisms of aging and the long-observed but mechanistically unexplained health benefits of taurine and spermidine.

## Introduction

Protein damage is a fundamental driver of aging and metabolic diseases (*1, 2*) Emerging evidence highlights carbon stress—the accumulation of damage from reactive acyl species (RAS) (*3*–*5*)—as a critical factor in these pathologies. Because RAS, most notably acyl-CoAs, contain electrophilic thioester moieties capable of non-enzymatic lysine acylation (*6*–*8*), their overproduction under carbon stress conditions, such as a high-fat diet, can markedly accelerate RAS-mediated protein damage (*9*–*11*). This is further exacerbated under metabolic stress, where the promiscuous activity of enzymes such as the histone acetyltransferase p300 reported as “acetyl spray”, leads to excessive protein acylation (*12, 13*). This aberrant acylation recently termed “spray acylation” disrupts protein function, perturbs the epigenomic landscape, ultimately accelerating age-associated declines (*3, 7, 8, 14, 15*) (**Fig. 1A**). Despite the prevalence of RAS-mediated damage, the endogenous metabolites responsible for scavenging these reactive species remain unidentified.

**Fig. 1.**
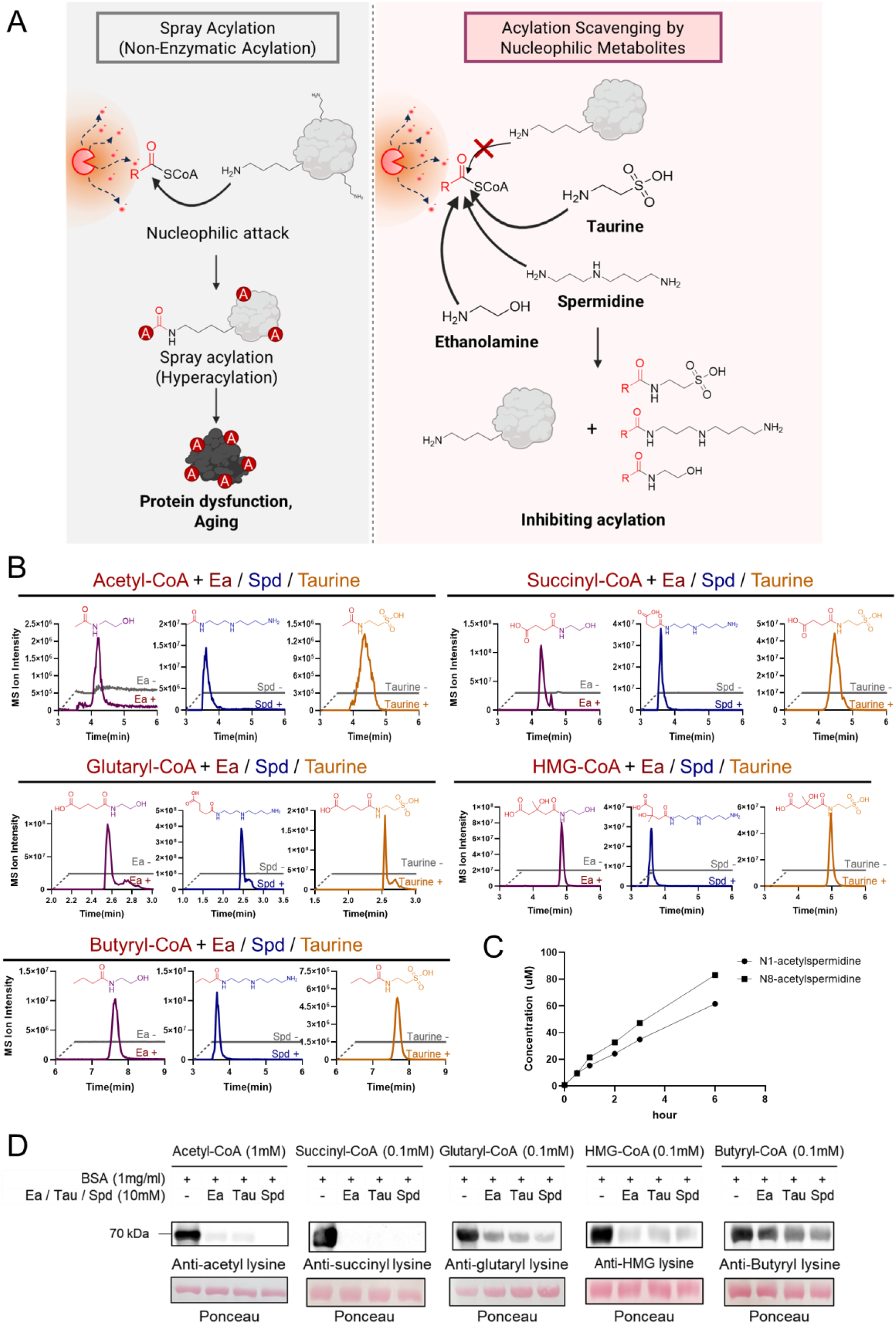
Nucleophilic metabolites scavenge acyl-CoAs to prevent non-enzymatic protein acylation. (A) Schematic illustrating the protective role of nucleophilic metabolites (taurine, spermidine, and ethanolamine) against acyl-CoA-mediated hyperacylation and consequent protein dysfunction. (B) Extracted ion chromatograms (EICs) of N-acylated TSEs. Indicated acyl-CoAs (acetyl-, succinyl-, glutaryl-, HMG-, and butyryl-CoA) were incubated in the presence or absence of scavengers (10 mM each of ethanolamine [Ea], taurine [Tau], or spermidine [Spd]) at pH 8.0, 37 °C for 16 h. (C) Time-course quantification of N1- and N8-acetylspermidine formation from an *in vitro* reaction of 1 mM acetyl-CoA and 10 mM spermidine (pH 8.0, 37 °C) over 6 h. (D) Immunoblot analysis of BSA acylation. BSA was incubated with indicated acyl-CoAs and TSE (10 mM each) at pH 8.0 and 37°C for 16 h.

Here, we identify taurine, spermidine, and ethanolamine (TSE) as endogenous small nucleophilic metabolites that scavenge reactive acyl species (RAS). While these molecules contain primary amine moieties capable of nucleophilic attack and have previously been associated with lifespan extension (*16*–*21*), the chemical basis for their metabolic effects remains poorly understood. We show that TSE react with freely diffusing acyl-CoAs and can also access and scavenge acyl intermediates that are bound within the active site of enzymes under physiological conditions. This ability to scavenge RAS within the enzyme pocket establishes a mechanism for limiting protein hyperacylation, a protective role physiologically demonstrated by spermidine’s direct interception of acetyl-CoA within the active site of the histone acetyltransferase p300.

We further examined the physiological relevance of this scavenging activity *in vivo*. In *Drosophila* overexpressing the p300 ortholog *nej*, spermidine-mediated prevention of hyperacetylation extended lifespan. In mice fed a high-fat diet, taurine supplementation led to the formation of N-fatty acyl taurines, indicating engagement of this scavenging chemistry in mammals. Together, these findings suggest a model in which endogenous nucleophilic metabolites mitigate carbon stress, thereby safeguarding cellular proteostasis.

### Taurine, spermidine, and ethanolamine scavenge acyl-CoAs and prevent protein acylation *in vitro*

To test whether taurine, spermidine, and ethanolamine (TSE) directly scavenge reactive acyl species (RAS), we examined their reactivity toward acyl-CoAs *in vitro* (**Fig. 1A**). A panel of acyl-CoAs—acetyl-CoA (1 mM), succinyl-CoA, glutaryl-CoA, HMG-CoA, and butyryl-CoA (0.1 mM)—was incubated with each of TSE at pH 8.0. Under these conditions, we detected the formation of N-acylated taurine, spermidine, and ethanolamine products (**Fig. 1B**). Tandem mass spectrometry confirmed characteristic fragment ions consistent with amide bond formation between the acyl moieties and TSE (**Extended Data Fig. 1, Table S1**), demonstrating direct nucleophilic scavenging of TSE. To further characterize the efficiency of these scavenging reactions, we performed a time-course kinetic analysis of spermidine acetylation, focusing on the formation of N1- and N8-acetylspermidine (**Fig. 1C**). When 1 mM acetyl-CoA was reacted with 10 mM spermidine, both products (N1- and N8-acetylspermidine) were detectable within 30 minutes, with their concentrations steadily increasing for 6 hours.

Notably, 10 mM of each TSE effectively prevented lysine acylation on BSA (**Fig. 1D**). This concentration is within the physiological range of taurine (5–50 mM) (*22, 23*) in the mitochondrial matrix, where acyl-CoAs are abundantly generated (*24, 25*). BSA acylation was further decreased in a dose-dependent manner with increasing TSE concentrations (50 μM–5 mM) (**Extended Data Fig. 2**). These data indicate that nucleophilic metabolites can rapidly intercept acyl-CoAs, effectively competing with protein lysine acylation.

We next examined whether TSE can also intercept reactive acyl intermediates bound within enzyme active sites, using the biotin ligase BirA and its engineered derivative TurboID as model systems. These enzymes generate a highly reactive biotinyl-AMP intermediate—chemically analogous to acyl-AMP. In Flp-In T-REx 293 cells stably expressing TurboID-V5-NES, co-treatment with taurine, spermidine, or ethanolamine during biotin labeling reduced global protein biotinylation in a dose-dependent manner without affecting TurboID expression (**Extended Data Fig. 3A–C**). LC-MS/MS analysis detected biotinyl-taurine, biotinyl-spermidine, and biotinyl-ethanolamine, confirming *in situ* nucleophilic scavenging of biotinyl-AMP (**Extended Data Fig. 3D**,**E**). Parallel *in vitro* reactions with recombinant wild-type BirA yielded the same biotinyl-TSE conjugates (**Extended Data Fig. 4**). Together, these results demonstrate that TSE can access and intercept reactive acyl-AMP intermediates within the catalytic pockets of “promiscuous” acyl-transfer enzymes—extending the scavenging mechanism beyond freely diffusing acyl-CoA capture.

### Polyamines block p300-mediated acetylation through nucleophilic scavenging

Given that TSE scavenged not only solution-phase acyl-CoAs but also active site-bound acyl intermediates in biochemical model systems, we investigated whether this mechanism could suppress aberrant protein acylation in physiologically relevant conditions. We focused on p300, recently characterized as an “acetyl spray” enzyme (*12, 26*), whose histone acetyltransferase (HAT) domain possesses a widely solvent-exposed substrate (i.e. acetyl-CoA) binding site (*27, 28*), suggesting high accessibility for proximal scavengers. Interestingly, although spermidine is a known inhibitor of p300 (*17, 29, 30*) that induces autophagy and promotes longevity (*17, 18, 21, 30, 31*), its precise molecular mechanism has remained elusive. We hypothesized that spermidine and other polyamines inhibit p300 by directly intercepting acetyl-CoA within the catalytic pocket.

To test this hypothesis, we overexpressed the p300 HAT domain in HEK293T cells and conducted untargeted metabolomic analysis **(Fig. 2A)**. Comparative analysis showed a profound increase in acetylated polyamines in p300-transfected cells relative to mock controls, including N8-acetylspermidine, N1,N8-diacetylspermidine, N-acetylputrescine, N1,N6-diacetylputrescine, N-acetylspermine, and N1,N12-diacetylspermine (**Fig. 2B**). Absolute quantification revealed substantial increases ranging from 32-to 732-fold for these species **(Fig. 2C)**. These metabolites were further validated through MS/MS spectral comparison with synthetic standards **(Fig. 2D)**. In parallel, the levels of free polyamines such as spermidine and spermine decreased by 1.97- and 3.21-fold, respectively, suggesting their consumption as endogenous substrates for p300-mediated reaction **(Extended Data Fig. 5)**.

**Fig. 2.**
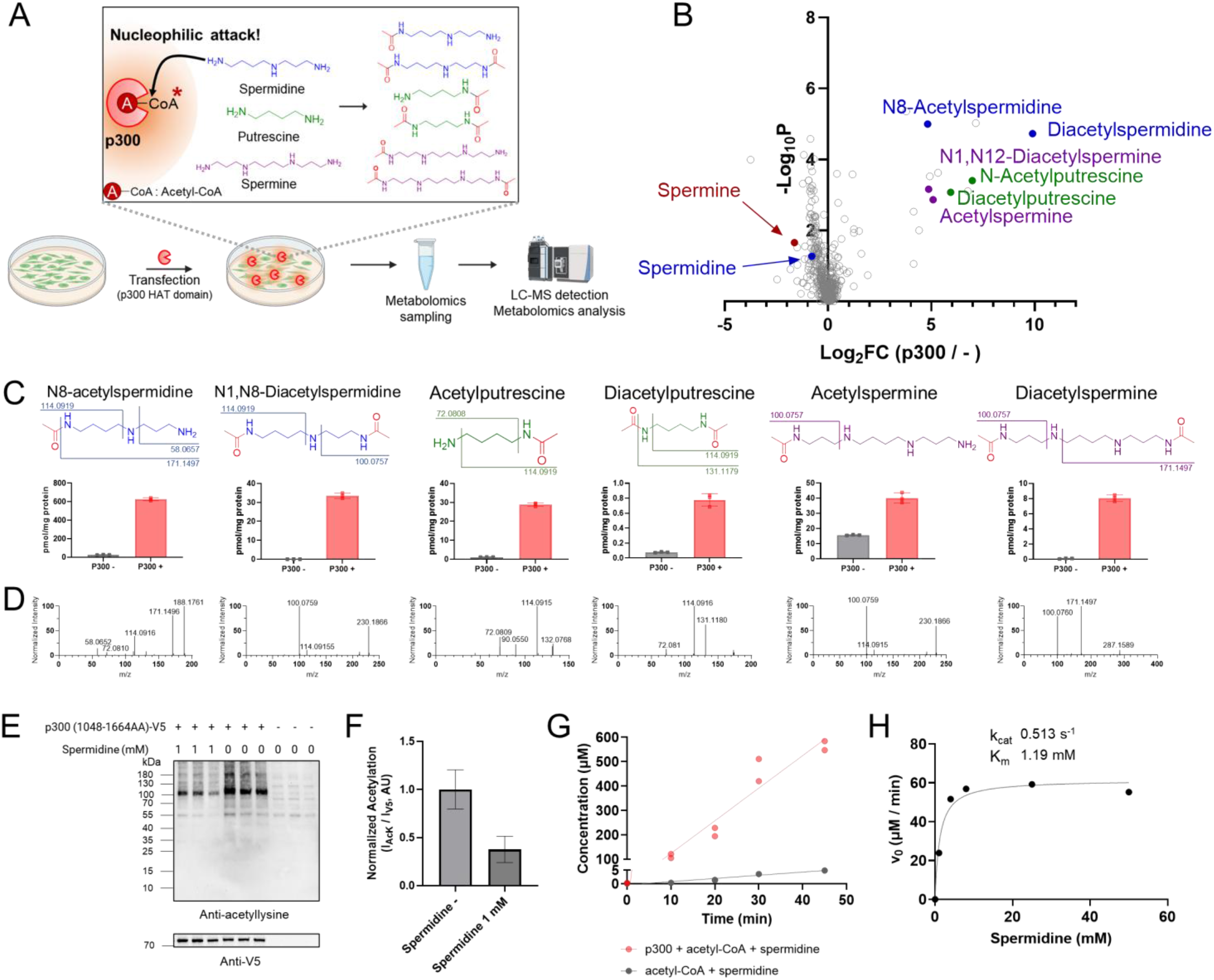
Spermidine prevents p300-mediated acetylation via nucleophilic scavenging. (A) Experimental workflow for metabolomics analysis of HEK293T cells transfected with the V5-tagged p300 HAT domain (residues 1048–1664; hereafter p300-HAT-V5). (B) Volcano plot showing the differential metabolome of p300-HAT-V5-overexpressing vs. mock-transfected cells (n = 3). Acetylated polyamines are highlighted. Metabolites were putatively annotated by molecular formula and ChemSpider database matching. P-values were calculated using an unpaired two-tailed Welch’s t-test. (C) Chemical structures and intracellular levels of mono- and di-acetylated polyamines in p300-HAT-V5-overexpressing cells. (D) MS/MS spectra identifying the indicated acetylated polyamines shown in (C). (E) Immunoblot analysis of acetyllysine and V5 in p300-HAT-V5-transfected HEK293T cells treated with or without 1 mM spermidine for 18 h. (F) Quantification of global acetylation from (E), normalized to p300 expression (I_AcK_ / I_V5_). (G) *In vitro* quantification of enzymatic and non-enzymatic N8-acetylspermidine production over time. Purified p300 HAT domain (1048–1664 AA, 2 µM) was incubated with 1 mM acetyl-CoA and 5 mM spermidine. (H) Kinetic analysis of *in vitro* p300 HAT-mediated N8-acetylspermidine production with spermidine.

Notably, p300-induced global protein acetylation was significantly reduced by co-treatment with spermidine, spermine, or putrescine each **(Fig. 2E and F, Extended Data Fig. 6)**. This was further supported by the observation that exogenous polyamine (spermidine, putrescine, or spermine) supplementation led to the intracellular accumulation of their corresponding acetylated metabolites **(Extended Data Fig. 7)**. *In vitro* assays using the recombinant p300 HAT domain showed that the enzyme markedly catalyzes N8-acetylspermidine production compared to the non-enzymatic rate (**Fig. 2G**). This suggests that spermidine preferentially reacts with the activated acetyl-CoA within the p300 active site. *In vitro* kinetic analysis of p300-mediated spermidine acetylation yielded a K_m_ of 1.19 mM (**Fig. 2H**), which aligns well with the reported intracellular spermidine concentrations (∼mM) (*32, 33*). Furthermore, the observed k_cat_ of 0.51 s^-1^ is comparable to the reported catalytic rates of p300 toward its native histone substrates (0.2 to 1 s^-1^) (*34*). These findings collectively indicate that polyamines protect proteins from p300-mediated aberrant acetylation through the nucleophilic scavenging of acetyl-CoA within the enzyme’s active site.

### Spermidine rescues p300-driven toxicity and extends lifespan in *Drosophila* through *in vivo* acetylation scavenging

To assess whether spermidine-mediated acetylation scavenging mitigates p300-driven toxicity in vivo, we employed a Drosophila melanogaster model overexpressing *nej*, the fly ortholog of human p300 (**Fig. 3A**). We first investigated whether spermidine engages in acetylation scavenging in vivo under elevated Nej activity conditions. Targeted metabolomic analysis of whole-fly extracts revealed a significant increase in acetylated polyamines, including N8-acetylspermidine and N1,N8-acetylspermidine, upon *nej* overexpression. These levels were further elevated by spermidine supplementation (**Fig. 3B**), further validating their role as acetylation scavengers in vivo.

**Fig. 3.**
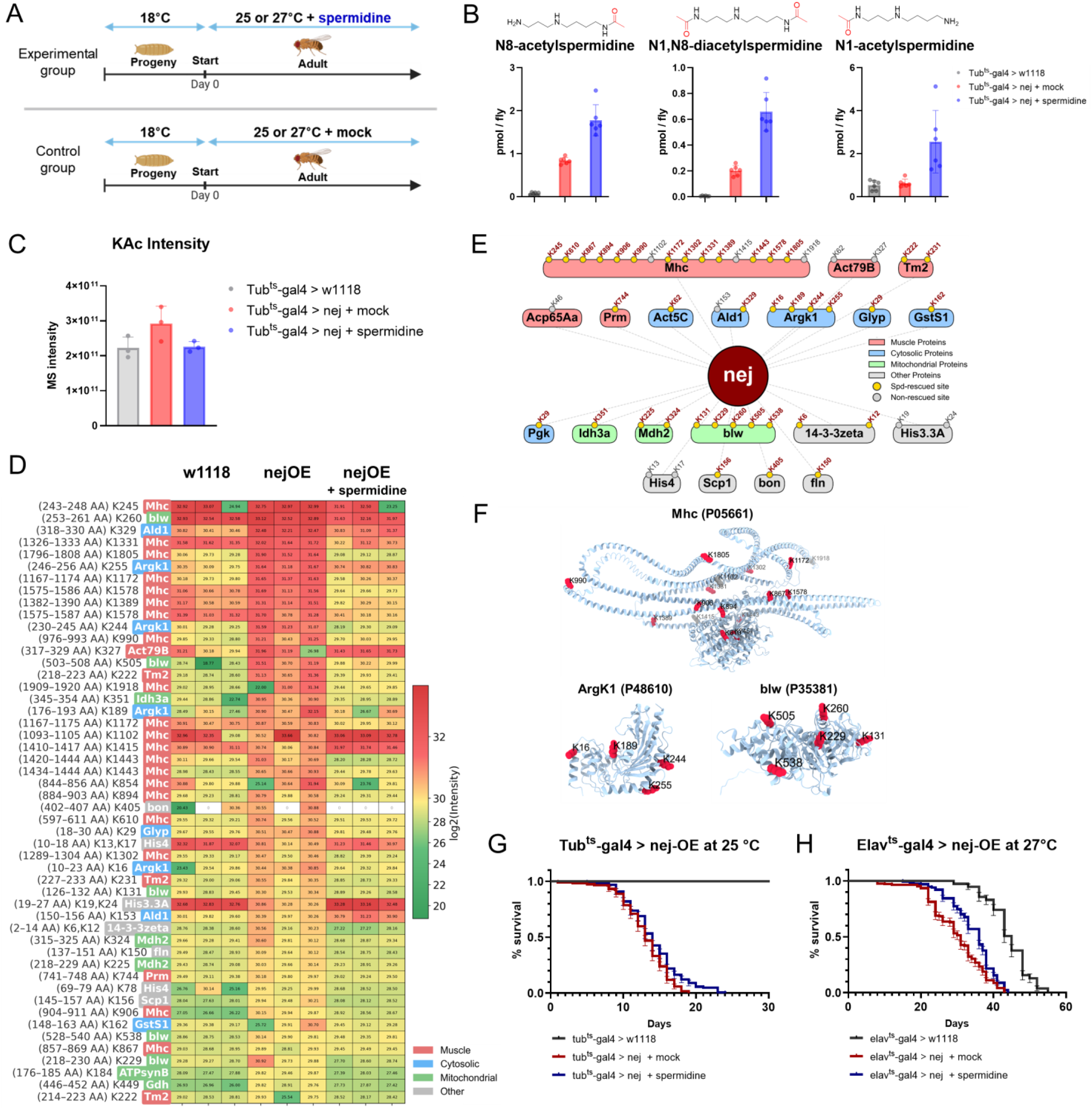
Nucleophilic scavenging of spermidine attenuates nej (p300)-mediated hyperacetylation and lifespan reduction in *Drosophila*. (A) Experimental scheme for *nej* overexpression using whole-body (tub-gal80ts;tubulin-Gal4) and neuron-specific (elav-gal4;;tub-gal80ts) drivers combined with spermidine treatment.(B) Levels of acetylated spermidines in whole-body *nej*-OE flies (N8-acetylspermidine, N1,N8-diacetylspermidine, and N1-acetylspermidine). (C) Comparison of lysine acetylation (KAc) intensities. (D) Heatmap of log2-transformed MS intensities for the top 50 Nej-dependent acetylated peptides. Rows represent individual acetylation sites labeled with gene name, peptide position range (amino acids), and modified lysine residues. (E) Acetylation network map of Nej target proteins. Gold circles indicate acetylation sites rescued by spermidine treatment. (F) AlphaFold-predicted three-dimensional structures of Mhc (P05661), Argk1 (P48610), and blw (P35381). Rescued sites by spermidine treatment are shown in red, while non-rescued sites are depicted as gray. (G, H) Survival curves of (G) whole-body and (H) neuron-specific *nej*-overexpressing flies with and without 5 mM spermidine treatment. Statistical significance was determined by log-rank (Mantel-Cox) test. Data are mean ± SD (n=3).

We then examined whether this spermidine-mediated scavenging influences the global acetylation landscape. Quantitative proteomics analysis of whole-body fly lysates revealed that *nej* overexpression (*nej*-OE) increased acetylated peptide MS intensities, which were markedly attenuated by spermidine treatment (**Fig. 3C**). To identify specific Nej-dependent acetylation targets that are scavenged by spermidine, we filtered our quantitative acetylproteomics dataset for peptides exhibiting hyperacetylation upon *nej* overexpression and subsequent reduction following spermidine treatment. A heatmap of the top 50 acetylated peptides ranked by MS intensity revealed distinct clusters of hyperacetylation in *nej*-overexpressing flies, which were profoundly restored following spermidine supplementation (**Fig. 3D**). Importantly, global protein levels of the top acetylation target proteins remained stable across all conditions (**Data S4**), confirming that the observed hyperacetylation reflects increased protein acetylation rather than changes in protein abundance. We subsequently mapped these top candidates into three principal categories: muscle, cytosolic, and mitochondrial metabolism. This landscape was further visualized using a Nej acetylation network map (**Fig. 3E**). Strikingly, dietary spermidine successfully mitigated Nej-induced hyperacetylation on the vast majority of these targeted lysine residues, confirming its role as an in vivo acetylation scavenger.

Detailed functional grouping highlighted muscle-associated proteins as one of the most severely impacted targets. Myosin heavy chain (Mhc), the core motor protein for muscle contraction, contained the highest number of hyperacetylated sites (16 sites observed within the top 50 targets). AlphaFold-based three-dimensional structural mapping showed that these hyperacetylated sites span critical domains of Mhc (**Fig. 3F**), suggesting that Nej-mediated aberrant acetylation globally compromises sarcomeric integrity and actin-myosin cross-bridge cycling. This aligns with vertebrate models where hyperacetylation of sarcomeric proteins—driven by stress-induced HATs (p300/PCAF) or inhibition of sarcomere-associated HDACs (HDAC3/4)—deregulates actin-myosin cross-bridge kinetics and directly contributes to cardiac and muscular dysfunction (*35, 36*). Spermidine treatment efficiently rescued 13 out of 16 mapped acetylation sites on Mhc, suggesting that spermidine directly preserves the structural and functional integrity of the muscle. Notably, spermidine treatment protected other critical structural components, including Tropomyosin (Tm2), Paramyosin (Parm), and non-muscle Myosin light chain (Mlp60A), from Nej-induced hyperacetylation (**Fig. 3D, E**).

Alongside muscle deterioration, *nej* overexpression induced widespread hyperacetylation of essential metabolic and mitochondrial enzymes. In the cytosol, Arginine kinase (Argk1), a key enzyme for rapid ATP buffering, displayed comprehensive hyperacetylation affecting its structural domains (**Fig. 3F**). Other critical cytosolic metabolic enzymes, such as Pgk, Ald1, GlyP, Act5C, and the antioxidant enzyme GstS1, were similarly hyperacetylated. Within the mitochondria, we observed extensive acetylation of the ATP synthase α subunit (blw) across multiple lysine sites in the *nej*-OE group **(Fig. 3D, F**), along with TCA cycle-associated enzymes including Idh3a, Mdh2, and Gdh, suggesting a critical impairment of bioenergetics and antioxidant defenses (*37*–*40*). Spermidine supplementation reverted the hyperacetylation of all mapped residues across these cytosolic and mitochondrial metabolic networks, effectively preserving mitochondrial bioenergetic machinery and cellular redox balance.

Consistent with prior studies linking p300 to cellular toxicity and senescence (*41*), whole-body *nej* overexpression resulted in a pronounced reduction in lifespan, indicative of deleterious hyperacetylation. Notably, dietary supplementation with 5 mM spermidine slightly improved survival in whole body *nej*-overexpressing flies (**Fig. 3G**). More significant lifespan extension was observed in flies with brain-specific *nej* overexpression upon spermidine supplementation (**Fig. 3H**). Feeding assays confirmed that these effects were not attributable to altered food intake (**Extended Data Fig. 8**), supporting a protective role of spermidine independent of caloric intake.

In conclusion, our data propose that spermidine prevents p300/Nej-mediated hyperacetylation by intercepting acetyl-CoAs through nucleophilic attack. By counteracting Nej-induced hyperacetylation across the muscle, cytosolic metabolism, TCA cycle and OXPHOS, spermidine protects them from proteotoxic collapse. The rescue of functionally distinct axes provides a compelling molecular basis for spermidine’s role as an endogenous scavenger that counteracts carbon stress in vivo.

### Taurine scavenges fatty acyl-CoAs and mitigates diet-induced obesity in mice

To evaluate reactive acyl species (RAS) scavenging in a physiological setting, we next focused on taurine in an obesity model. Previous studies have shown that dietary taurine supplementation attenuates high-fat diet (HFD)– induced weight gain in mice (*40*–*45*), yet the underlying metabolic basis remains unclear. To explore whether this phenotype involves RAS scavenging, we examined taurine metabolism under HFD conditions, which are known to promote the accumulation of fatty acyl-CoAs and lipotoxicity (*46*) (**Fig. 4A**). C57BL/6N mice were fed an HFD for three weeks with or without taurine supplementation (5% w/v in drinking water). Consistent with prior reports (*40*–*45*), taurine supplementation significantly attenuated HFD-induced body weight gain compared with vehicle-treated controls (**Fig. 4B**). While HFD feeding modestly reduced endogenous taurine levels, supplementation restored its abundance, confirming effective systemic delivery (**Extended Data Fig. 9**) (*40*).

**Fig. 4.**
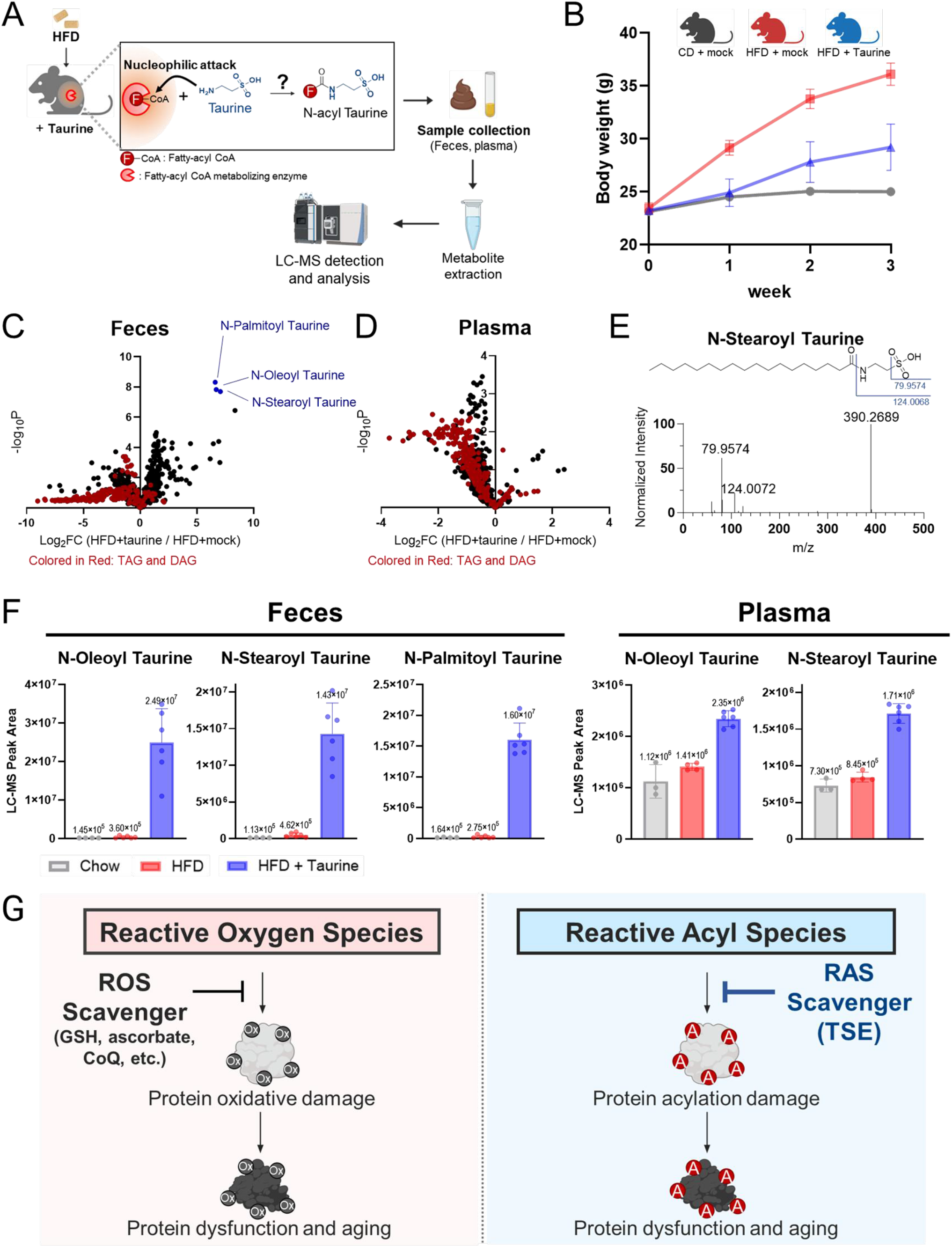
Taurine supplementation promotes N-fatty acyl taurine formation by intercepting fatty acyl-CoAs in high-fat diet–fed mice. (A) Experimental scheme validating the in vivo scavenging reaction between fatty acyl-CoA and taurine. Plasma and fecal samples were analyzed by LC-MS/MS to detect N-fatty acyl taurine. (B) Body weight changes in C57BL/6N mice fed a high-fat diet (HFD) with (n=6) or without (n=6) taurine supplementation (5% w/v in drinking water), compared to chow diet controls (n=4). (C, D) Volcano plots of the plasma and (D) fecal lipidomes comparing taurine-treated vs. mock-treated HFD mice. (E) Fragmentation pattern and MS/MS spectrum of N-stearoyl taurine. (F) Levels of (left) oleoyl-, stearoyl-, and palmitoyl-taurine in feces, and (right) oleoyl- and stearoyl-taurine in plasma collected via submandibular (cheek) bleed. Data are mean ± SD; n = 3; ^*^P < 0.05, unpaired two-tailed Welch’s t-test. (G) Conceptual model comparing the reactive oxygen species (ROS) defense system with the reactive acyl species (RAS) defense system.

To determine whether taurine directly reacts with fatty acyl-CoAs in vivo, we performed untargeted lipidomic analyses of plasma and fecal samples from HFD-fed mice with or without taurine supplementation. Taurine supplementation was associated with a broad remodeling of the lipidome in both samples, including an overall reduction in lipid abundance that correlated with the observed attenuation of weight gain (**Fig. 4C and D**). Most notably, we observed a marked accumulation of N-fatty acyl taurines (NATs), which were identified by a diagnostic fragment ion characteristic of N-taurinylation (**Fig. 4E**). In fecal samples, levels of N-stearoyl-, N-oleoyl-, and N-palmitoyl-taurine increased by approximately 30-to 70-fold following taurine supplementation. Consistent with this, circulating levels of N-oleoyl- and N-stearoyl-taurine in plasma were also elevated (**Fig. 4F**). Together, these data provide direct in vivo evidence that dietary taurine directly intercepts reactive fatty acyl-CoAs to form stable N-fatty acyl taurine conjugates, consistent with nucleophilic scavenging of diet-derived RAS.

## Discussion

Taurine, spermidine, and ethanolamine are well-known contributors to metabolic health and longevity across species, yet their beneficial effects have been attributed to diverse pathways—ranging from tRNA modification (*20*) and autophagy (*18, 21, 29, 31*) to lipid homeostasis (*19*). Despite extensive study, a unifying molecular principle for these effects has remained elusive. Here, we propose that nucleophilicity is the shared chemical property underlying their common protective function. By directly intercepting RAS, TSE limit aberrant protein acylation and protect the proteome from carbon stress. This nucleophilic scavenging mechanism provides a chemical framework that integrates the pleiotropic biological effects of TSEs into a unified model of metabolic defense.

Protein acylation is traditionally viewed as a balance between acyltransferases and deacylating enzymes such as sirtuins (*24, 25*). Our findings add a chemically distinct layer of regulation in which TSE intercept reactive acyl donors upstream of protein modification. This interception occurs both in solution and within specific catalytic environments, such as the active site of p300. The scavenging within the catalytic pocket suggests that the efficacy of chemical scavenging depends on the structural accessibility and microenvironment of the enzyme’s catalytic pocket. Consequently, this mechanism is unlikely to apply uniformly; instead, TSE-mediated scavenging may preferentially impact enzymes with high solvent exposure and broad acyl-donor specificity, while sparing tightly regulated reactions essential for cellular function. This selectivity explains why systemic TSE supplementation extends healthspan without inducing toxicity.

This mechanism is exemplified by the reaction between taurine and diet-derived fatty acyl-CoAs under high-fat diet conditions in mammals. The resulting N-fatty acyl taurines act as metabolic dead-end products that are efficiently eliminated via fecal excretion (**Fig. 4F**). This pathway mirrors classical drug detoxification, where drug-taurine conjugation facilitates their removal rather than reintegration into metabolism (*47, 48*). By converting reactive fatty acyl-CoAs into inert, excretable conjugates, taurine reduces acyl-donor pressure without perturbing systemic carbon flux. Consequently, as TSE are not readily funneled into central carbon metabolism, their exogenous intake does not impose the energetic or metabolic flux burdens associated with the high-level consumption of other nucleophilic amine metabolites, such as amino acids.

Our findings unveil a previously unrecognized biosynthetic pathway for acyl-TSE species, demonstrating their formation non-enzymatically or within the active site of “acyl-spray” enzymes such as p300 (*12, 13*). This mechanism contrasts with their traditional classification as products primarily formed via dedicated enzymatic processes (*49*). As terminal products of this scavenging chemistry, N-acyl-TSE species could serve as biomarkers of *in vivo* carbon stress, reflecting cumulative burden of aberrant acylation. Although these metabolites are often overlooked in standard metabolomic panels, their mass spectrometric signatures established in this study (**Extended Data Fig. 1**) enable their identification and inclusion in future metabolomic profiling.

It is worth noting that the observed scavenging by TSE is not universally applicable to all RAS-generating enzymes. The efficacy appears to depend on specific enzyme-scavenger pairings (e.g. p300-spermidine), likely governed by the micro-environment and accessibility of the enzyme active site. However, this selectivity suggests that a diverse pool of endogenous nucleophilic metabolites may be required to scavenge distinct enzymatic reactions. Thus, while this study focuses on three specific metabolites (TSE), systematically identifying other endogenous nucleophiles that scavenge enzymatic reactions will be essential for fully elucidating their broader metabolic significance.

In summary, our study identifies nucleophilic scavenging by endogenous metabolites, taurine, spermidine, and ethanolamine (TSE) as a fundamental chemical defense against protein acylation damage. This mechanism directly parallels the role of classic antioxidant scavengers (e.g., vitamin C, glutathione, coenzyme Q) against ROS stress (*50, 51*), positioning TSE as intrinsic scavengers against reactive acyl species (RAS). Just as cells rely on antioxidant scavengers such as glutathione and ascorbate to neutralize reactive oxygen species (ROS), our data reveal that TSE constitute a parallel scavenging system that neutralizes RAS and thereby protects the proteome from carbon stress (**Fig. 4G**). Collectively, these findings establish nucleophilic scavenging of RAS as a new paradigm with broad implications for combating aging, metabolic disease, and carbon stress.

## Acknowledgments

We are grateful to Dr. Richard Binari and Dr. Myeonghoon Han for assisting with the fly experiments, Minsang Hwang for the acetylome analysis, Jae-Yoon Jo for the preparation of in-house packed capillary column, and Chang-Mo Yoo for valuable discussions.

## Author contributions

Conceptualization: MKL, HWR

Data curation: MKL, SMK, SHL

Formal analysis: MKL, SMK, SHL

Investigation: MKL, SMK, SHL, JHY, CRC, MT

Supervision: JSK, NP, YPK, HWR

Writing – original draft: MKL, HWR

Writing – review & editing: JSK, NP, YPK, HWR

## Data, code, and materials availability

The authors declare that all data supporting the findings of this study are available in the paper and Supplementary Information. Additional datasets related to this study are available from the corresponding authors on reasonable request. Our raw mass analysis files were deposited to the Proteo-meXchange (http://proteomecentral.proteomexchange.org) via the PRIDE partner repository with identifier PXD033869 (Username: / Password: r3hgUtUM).

## Extended Data and Supplementary Information

Methods

Extended Data Figs. 1 to S9

Tables S1 to S3

Data S1 to S6

## References

1. C. López-Otín, M. A. Blasco, L. Partridge, M. Serrano, G. Kroemer, Hallmarks of aging: An expanding universe. Cell 186, 243–278 (2023).

2. M. S. Hipp, The proteostasis network and its decline in ageing. Nat. Rev. Mol. Cell Biol. 20, 421–435 (2019).

3. A. G. Trub, M. D. Hirschey, Reactive Acyl-CoA Species Modify Proteins and Induce Carbon Stress. Trends Biochem. Sci. 43, 369–379 (2018).

4. G. R. Wagner, M. D. Hirschey, Nonenzymatic Protein Acylation as a Carbon Stress Regulated by Sirtuin Deacylases. Mol. Cell 54, 5–16 (2014).

5. B. Zhang, F. C. Schroeder, Mechanisms of metabolism-coupled protein modifications. Nat. Chem. Biol. 21, 819– 830 (2025).

6. G. R. Wagner, R. M. Payne, Widespread and enzyme-independent Nε-acetylation and Nε-succinylation of proteins in the chemical conditions of the mitochondrial matrix. Journal of Biological Chemistry 288, 29036–29045 (2013).

7. S. Softic, et al., Dietary Sugars Alter Hepatic Fatty Acid Oxidation via Transcriptional and Post-translational Modifications of Mitochondrial Proteins. Cell Metab. 30, 735-753.e4 (2019).

8. A. A. Kendrick, et al., Fatty liver is associated with reduced SIRT3 activity and mitochondrial protein hyperacetylation. Biochemical Journal 433, 505–514 (2011).

9. P. Gut, et al., SUCLA2 mutations cause global protein succinylation contributing to the pathomechanism of a hereditary mitochondrial disease. Nat. Commun. 11, 1–14 (2020).

10. B. T. Weinert, et al., Time-Resolved Analysis Reveals Rapid Dynamics and Broad Scope of the CBP/p300 Acetylome. Cell 174, 231-244.e12 (2018).

11. B. M. Dancy, P. A. Cole, Protein lysine acetylation by p300/CBP. Chem. Rev. 115, 2419–2452 (2015).

12. B. R. Sabari, D. Zhang, C. D. Allis, Y. Zhao, Metabolic regulation of gene expression through histone acylations. Nat. Rev. Mol. Cell Biol. 18, 90–101 (2017).

13. C. Choudhary, B. T. Weinert, Y. Nishida, E. Verdin, M. Mann, The growing landscape of lysine acetylation links metabolism and cell signalling. Nat. Rev. Mol. Cell Biol. 15, 536–550 (2014).

14. P. Singh, et al., Taurine deficiency as a driver of aging. Science. 380, eabn9257 (2023).

15. F. Madeo, T. Eisenberg, F. Pietrocola, G. Kroemer, Spermidine in health and disease. Science. 359, eaan2788 (2018).

16. S. J. Hofer, et al., Spermidine Is Essential for Fasting-Mediated Autophagy and Longevity. Nature Cell Biology. 26, 1571–1584 (2024).

17. T. Eisenberg, et al., Cardioprotection and lifespan extension by the natural polyamine spermidine. Nat. Med. 22, 1428–1438 (2016).

18. P. Rockenfeller, et al., Phosphatidylethanolamine positively regulates autophagy and longevity. Cell Death Differ.22, 499–508 (2015).

19. P. Rockenfeller, D. Carmona-gutierrez, F. Pietrocola, G. Kroemer, F. Madeo, Ethanolamine : A novel anti-aging agent. Mol. Cell. Oncol., 3, e1019023 (2016).

20. C. J. Jong, P. Sandal, S. W. Schaffer, The role of taurine in mitochondria health: More than just an antioxidant. Molecules 26, 1–21 (2021).

21. JG Jacobsen, LH Smith, Biochemistry and Physiology of Taurine and Taurine Derivatives. Physiol. Rev. 48, 424– 511 (1968).

22. A. E. Ringel, S. A. Tucker, M. C. Haigis, Chemical and Physiological Features of Mitochondrial Acylation. Mol. Cell 72, 610–624 (2018).

23. C. Carrico, J. G. Meyer, W. He, B. W. Gibson, E. Verdin, The Mitochondrial Acylome Emerges: Proteomics, Regulation by Sirtuins, and Metabolic and Disease Implications. Cell Metab. 27, 497–512 (2018).

24. Y. B. Lee, H. W. Rhee, Spray-type modifications: an emerging paradigm in post-translational modifications. Trends Biochem. Sci. 49, 208–223 (2024).

25. J. Maksimoska, D. Segura-Peña, P. A. Cole, R. Marmorstein, Structure of the p300 Histone Acetyltransferase Bound to Acetyl-Coenzyme A and Its Analogues. Biochemistry 53, 3415–3422 (2014).

26. X. Liu, et al., The structural basis of protein acetylation by the p300/CBP transcriptional coactivator. Nature 451, 846–850 (2008).

27. F. Pietrocola, et al., Spermidine induces autophagy by inhibiting the acetyltransferase EP300. Cell Death Differ. 22, 509–516 (2015).

28. S. J. Hofer, et al., Mechanisms of spermidine-induced autophagy and geroprotection. Nat. Aging 2, 1112–1129 (2022).

29. T. Eisenberg, et al., Induction of autophagy by spermidine promotes longevity. Nat. Cell Biol. 11, 1305–1314 (2009).

30. A. E. Pegg, Functions of Polyamines in Mammals. Journal of Biological Chemistry 291, 14904–14912 (2016).

31. C. Biology, K. Igarashi, K. Kashiwagi, Modulation of cellular function by polyamines. 42, 39–51 (2010).

32. P. R. Thompson, et al., Regulation of the p300 HAT domain via a novel activation loop. Nat. Struct. Mol. Biol. 11, 308–315 (2004).

33. M. P. Gupta, S. A. Samant, S. H. Smith, S. G. Shroff, HDAC4 and PCAF Bind to Cardiac Sarcomeres and Play a Role in Regulating Myofilament Contractile Activity. Journal of Biological Chemistry 283, 10135–10146 (2008).

34. S. A. Samant, V. B. Pillai, N. R. Sundaresan, S. G. Shroff, M. P. Gupta, Histone Deacetylase 3 (HDAC3)-dependent Reversible Lysine Acetylation of Cardiac Myosin Heavy Chain Isoforms Modulates Their Enzymatic and Motor Activity. Journal of Biological Chemistry 290, 15559–15569 (2015).

35. B.-H. Ahn, et al., A role for the mitochondrial deacetylase Sirt3 in regulating energy homeostasis. Proceedings of the National Academy of Sciences 105, 14447–14452 (2008).

36. S. Someya, et al., Sirt3 Mediates Reduction of Oxidative Damage and Prevention of Age-Related Hearing Loss under Caloric Restriction. Cell 143, 802–812 (2010).

37. M. D. Hirschey, et al., SIRT3 regulates mitochondrial fatty-acid oxidation by reversible enzyme deacetylation. Nature 464, 121–125 (2010).

38. S. Zhao, et al., Regulation of Cellular Metabolism by Protein Lysine Acetylation. Science 327, 1000–1004 (2010).

39. P. Sen, et al., Histone Acetyltransferase p300 Induces De Novo Super-Enhancers to Drive Cellular Senescence. Mol. Cell 73, 684-698.e8 (2019).

40. N. Tsuboyama-Kasaoka, et al., Taurine (2-Aminoethanesulfonic Acid) deficiency creates a vicious circle promoting obesity. Endocrinology 147, 3276–3284 (2006).

41. T. M. Batista, et al., Taurine supplementation improves liver glucose control in normal protein and malnourished mice fed a high-fat diet. Mol. Nutr. Food Res. 57, 423–434 (2013).

42. J. C. Santos, et al., Taurine supplementation ameliorates glucose homeostasis, prevents insulin and glucagon hypersecretion, and controls β, α, and δ-cell masses in genetic obese mice. Amino Acids 47, 1533–1548 (2015).

43. Y. Guo, B. Li, W. Peng, L. Guo, Q. Tang, Taurine-mediated browning of white adipose tissue is involved in its antiobesity effect in mice. Journal of Biological Chemistry 294, 15014–15024 (2019).

44. I. White, F. Tissue, Taurine Stimulates Thermoregulatory Genes in Brown Fat Tissue and Muscle without an Influence on Obese Mouse Model. Foods 9, 1–14 (2020).

45. A. D. Karikkakkavil Prakashan, S. P. Muthukumar, A. Martin, Taurine Activates SIRT1/AMPK/FOXO1 Signaling Pathways to Favorably Regulate Lipid Metabolism in C57BL6 Obese Mice. Mol. Nutr. Food Res. 68, 1–11 (2024).

46. Y. Du, et al., Lysine Malonylation Is Elevated in Type 2 Diabetic Mouse Models and Enriched in Metabolic Associated Proteins. Molecular and Cellular Proteomics 14, 227–236 (2015).

47. K. M. Knights, M. J. Sykes, J. O. Miners, Amino acid conjugation: contribution to the metabolism and toxicity of xenobiotic carboxylic acids. Expert Opin. Drug Metab. Toxicol. 3, 159–168 (2007).

48. M. Darnell, L. Weidolf, Metabolism of Xenobiotic Carboxylic Acids: Focus on Coenzyme A Conjugation, Reactivity, and Interference with Lipid Metabolism. Chem. Res. Toxicol. 26, 1139−1155 (2013).

49. J. Z. Long, et al., The Secreted Enzyme PM20D1 Regulates Lipidated Amino Acid Uncouplers of Mitochondria. Cell 166, 424–435 (2016).

50. H. Sies, Redox Biology Oxidative stress : a concept in redox biology and medicine. Redox Biol. 4, 180–183 (2015).

51. T. Finkel, N. J. Holbrook, Oxidants, oxidative stress and the biology of ageing. Nature 408, 239–247 (2000).

